# LARP1 regulates metabolism and mTORC1 activity in cancer

**DOI:** 10.1101/2022.09.04.506559

**Authors:** James Chettle, Zinaida Dedeic, Roman Fischer, Iolanda Vendrell, Leticia Campo, Alistair Easton, Molly Browne, Josephine Morris, Hagen Schwenzer, Pauline Lascaux, Rik Gijsbers, Elisabete Pires, Daniel J. Royston, David J. P. Ferguson, An Coosemans, Benedikt Kessler, James McCullagh, Ahmed A. Ahmed, Kristijan Ramadan, Martin Bushell, Adrian L. Harris, Colin R. Goding, Sarah P. Blagden

**Author notes:** Corresponding authors James Chettle, Department of Oncology, Roosevelt Drive, University of Oxford, OX3 7DQ, UK, Sarah Blagden, Department of Oncology, Roosevelt Drive, University of Oxford, OX3 7DQ, UK.

## Abstract

The protein mammalian target of rapamycin (mTOR) is a master regulator of cell homeostasis. Although mTOR is aberrantly overactivated in 70% ovarian cancers, mTOR cascade inhibitors (such as those blocking the kinase activity of mTOR itself or upstream kinases PI3K/AKT) have demonstrated disappointing activity in ovarian cancer clinical trials. These findings indicate that, despite its pivotal role in normal cells, hyperactivated mTOR does not act as a master regulator of metabolism in this cancer context. Surprisingly, we have identified that the RNA binding protein LARP1, a known phospho-target of mTORC1 and activator of ribosomal biogenesis, is responsible for metabolic reprogramming in mTOR-dysregulated cancers. LARP1 post-transcriptionally regulates the expression of several hundred rate-limiting enzymes involved in multiple aspects of metabolism, including glycolysis and oxidative phosphorylation. Through this mechanism LARP1 sustains ATP production and mTORC1 localisation on the lysosome, thereby activating cell proliferation despite the scarcity of extracellular nutrients. Our findings show that, by sustaining global cellular metabolism in response to growth factor signalling, LARP1 has a central post-transcriptional role in controlling mTORC1 localisation and driving cancer progression, a key cancer hallmark.

## Introduction

In normal cells, the serine/threonine kinase mammalian target of rapamycin (mTOR) is known as a ‘master regulator’ of nutrient-sensitive growth regulation via its two effector complexes mTORC1 and mTORC2^1–3^. Multiple studies have demonstrated that mTORC1 integrates intracellular and extracellular cues including from growth factors, stress signals, and the availability of ATP, oxygen and amino acids, to control anabolic processes including protein and lipid synthesis, cell growth and cell cycle progression^4,5^. Upon growth factor stimulation and/or heightened availability of ATP and amino acids, mTORC1 is activated by relocating to the surface of the lysosome where it rapidly increases protein synthesis to fuel anabolism^6^. To achieve this, lysosomal mTORC1 phosphorylates translation initiation proteins S6K and eIF4E-binding protein (4E-BP)^7,8^. Once phosphorylated, 4E-BP activates 5’cap-dependent protein synthesis by releasing its inhibitory hold on eIF4E, freeing eIF4E to bind to the 5’cap of mRNA along with other components of the eIF4F complex^9^. mTOR simultaneously supports ribosome availability by increasing ribosomal biogenesis through the RNA binding protein (RBP) LARP1^10–13^. Upon mTORC1 phosphorylation, LARP1 releases its inhibitory hold on ribosomal mRNAs, all of which carry terminal oligopyrimidine (TOP) motifs immediately adjacent to their 5’ cap (they are termed TOP mRNAs)^9,12,14^. Release of phosphorylated LARP1 from these TOP motifs frees their 5’ caps to bind eIF4E, assemble eIF4F complexes and activate coordinated synthesis of the ribosomal machinery required for translation of cell growth and proliferation^15,16^. At the same time as activating protein synthesis, mTORC1 phosphorylates other targets such as TFEB, MTHFD2, SREBP1 and HIF1α to drive lysosome biogenesis, nucleotide synthesis, lipid metabolism, glycolysis and mitochondrial oxidative metabolism^17,18^. The net result of mTOR activation is thus the ready availability of anabolic macromolecules and machinery to fuel cell proliferation^19^.

In an estimated 30% all cancers, despite existing in hostile environmental conditions in which nutrients are in short supply, cellular mTOR-signalling is paradoxically hyperactive^20^. In the minority (approximately 3%), this is due to a direct *mTOR*-activating gene mutation but, for the majority, mTOR hyperactivation is secondary to permissive signalling from growth factors, mutations in upstream positive (such as PIK3CA or AKT) or negative regulators (PTEN), or in interacting pathway genes^1,21,22^. The assumption that, in cancer cells, hyperactive mTOR retains its function as a master anabolic regulator has prompted numerous clinical trials of mTOR inhibitors^23^. For example, in ovarian cancer, where mTOR hyperactivation is observed in 70% cases, clinical trials have been conducted using mTOR inhibitors alone or in combination with chemotherapy but have yielded disappointing results^24,25^. In mTOR hyperactive cancer cell lines, treatment with mTOR inhibitors causes only a temporary anti-proliferative effect but little impact on overall metabolism or viability, indicating that other factors lie upstream of mTOR in driving metabolism and promoting viability in these cancer cells^26^.

To explore this further we looked at the relationship between mTORC1 and its effector protein LARP1 in cancer. LARP1 is abundantly expressed across epithelial malignancies, including ovarian, lung, cervical and hepatocellular carcinoma.^27,28^ In patients with ovarian cancer, high tumour LARP1 expression is associated with worse progression-free and overall survival^29^. By contrast, in normal cells, LARP1 is expressed at low levels where it predominantly acts as a transducer of mTORC1 signalling and effector of ribosomal biogenesis^9,14,30^. Using cell lines and in ovarian tumours we demonstrate that LARP1 is crucial for driving high-level metabolism in mTOR-hyperactivated cancers. Furthermore, we show for the first time that LARP1 regulates mTORC1 localisation on the lysosome and can threreby potentiate mTORC1 activity. Unlike inhibition of mTOR that causes a transient anti-proliferative effect, depletion of LARP1 is lethal in these cancer cells. Therefore, through its role in driving metabolism, cell survival and cell proliferation (the latter via mTORC1 activation), LARP1 is a post-transcriptional master regulator of metabolism in mTOR-hyperactivated ovarian cancer.

## Materials & Methods

### Cell culture

OVCAR-8, U2OS and HeLa cells were kindly provided by the Ovarian Cancer Action Biobank at Imperial College and genotyped prior to use. SKOV3 and HEY-A8 cells were purchased from ATCC. Unless otherwise stated, cells were cultured in RPMI with 10% foetal bovine in the presence of penicillin (50 U/ml) and streptomycin (50 μg/ml). All cells were negative for *Mycoplasma* upon regular testing with the MycoAlert Mycoplasma Detection Kit (Lonza; LT07-318). LARP1 knockout cell line generation was performed using Edit-R^™^ CRISPR-Cas9 Gene Engineering, using All-in-one lentiviral sgRNA particles (Dharmacon, Source clone ID: VSGHSOM_28505936). After lentiviral transduction, cells were selected by growth in puromycin (10 μg/ml) and individuals clones were expanded and sequenced. Two clones derived from OVCAR-8 cells, clone 6 and clone 9, displayed editing across all three copies of the LARP1 gene on chromosome 5. Clone 8 was confirmed by sequencing to be genetically identical to wild-type cells.

### siRNA knockdown and LARP1 expression vector transfection

Knockdown of LARP1 by siRNA was performed using Lipofectamine RNAiMax (ThermoFisher; 13778075). The following siRNAs were used to target LARP1 (siLARP1-1: Sense:AGACUCAAGCCAGACAUCA-dTdT, Antisense:UGAUGUCUGGCUUGAGUCU-dTdT siLARP1-2: Sense:CAUGACUCUUGACAUCCUA-dTdT, Antisense:UAGGAUGUCAAGAGUCAUG-dGdA siLARP1-3: Sense:GCACCGAGCUUUACAGUGU-dTdT, Antisense:ACACUGUAAAGCUCGGUGC-dTdG). The following siRNA was used as a non-targeting control (Sense:GGUCCGGCUCCCCCAAAUG-dTdT Antisense:CAUUUGGGGGAGCCGGACC-dTdT). The LARP1 expression vector was prepared by cloning full length *LARP1* (encoding the 1096 amino acid isoform of LARP1, UniProtKB/Swiss-Prot: Q6PKG0.2) was cloned into the expression vector pcDNA4/HisMax A (ThermoFisher, V86420). Transfection was performed with Fugene6 (Promega) using a 3:1 ratio of Fugene6 to plasmid, and 2 μg plasmid per well of a six well plate.

### mRNA and protein quantitation

mRNA levels of individual genes were measured by quantitative RT-PCR, normalised to levels of *ACTB* mRNA. For quantitative RT-PCR, mRNA was extracted from cells using the GenElute mammalian Total RNA miniprep extraction kit (Merck, RTN70-1KT) and was reverse transcribed to synthesise cDNA using High Capacity cDNA Reverse Transcription Kit (ThermoFisher, 4368814). mRNA concentrations were equalised by nanodrop prior to reverse transcription. Quantitative PCR was performed using Fast SYBR^™^ green master mix (ThermoFisher, 4385610) on a StepOnePlus ^™^ Real-Time PCR system. Primer sequences used are shown in **Supplementary Table S4**. Western blots were imaged using an Odyssey scanner (Licor), with densitometry analysis performed using ImageStudio software (Licor). Antibodies used in western blotting are listed in **Supplementary Table S5**.

### Proteomics and metabolomics

Three 10 cm dish plates of OVCAR-8 cells were treated with non-targeting siRNA and three with siLARP1-2. Cells were harvested at 48h after siRNA transfection and lysed in RIPA buffer (25 mM TrisHCl pH 7.6, 150 mM NaCl, 1%NP-40, 1% sodium deoxycholate, 0.1% SDS). 150 ug of protein were precipitated in methanol:chloroform after reduction with 20mM DTT (30min) and alkylation with 40mM Iodoacetamide (30 min). Precipitates were solubilised in 8M urea (20mM HEPES, pH 8) and digested with Lysine C (1:50 ratio; Promega) for 4h at 37C in 6M urea followed by an overnight trypsin digestion (1:25 ratio; Promega) at 37C in 1M urea, stopped by adding 1% TFA (final concentration). Tryptic peptide mixtures were desalted using SOLA-HRP SPE columns (ThermoFisher), dried and reconstituted in LC-MS/MS water containing 2% acetonitrile, 0.1 %TFA. Approximately 200ng of tryptic peptides were analysed by liquid chromatography tandem mass spectrometry (LC-MS/MS) using Ultimate 3000 UHPLC (ThermoFisher) connected to an Orbitrap Fusion Lumos Tribrid (ThermoFisher). Eluted peptides were analysed on an Orbitrap Fusion Lumos Tribrid platform (instrument control software v3.3). Data were acquired in data-dependent mode, using the universal method as described previously with the advance peak detection (APD) enabled.^31^ Survey scans were acquired in the Orbitrap at 120 k resolution over a m/z range 400 -1500, AGC target of 4e5 and S-lens RF of 30. Fragment ion spectra (MS/MS) were obtained in the Ion trap (rapid scan mode) with a Quad isolation window of 1.6, 40% AGC target and a maximum injection time of 30 ms, with HCD activation and 28% collision energy. Raw mass spectrometry data were analysed using MaxQuant (v1.6.10.43), searched against the UniProt-Swissprot human database using the Andromeda data-search engine at 1% protein false discovery rate. Data were quantified using label free quantitation (LFQ) using the match between runs and selecting the iBAQ. The Maxquant protein group output was further analysed using Perseus (1.6.2.2). A two-sample student t-test was applied combined with a Permutation-FDR correction (5%). Proteins with a significant student t-test value (q<0.05) were selected, normalised using the Z score and hierarchically classified.

For the metabolomics analysis, nine 10 cm plates of OVCAR-8 cells were treated with a non-targeting siRNA and were treated with either siLARP1-1,2 or 3. Samples were harvested at 48 h after siRNA transfection by methanol precipitation and subjected to LC-MS and statistical analysis following a previously described method.^32^

### Oxygen consumption analysis by Seahorse Mito Stress® assay

2×10^4^ – 4×10^4^ cells per well were plated onto a Seahorse XFe96 FluxPak plate (Agilent, 102416-100). On the day of the assay, media was replaced with Seahorse XF DMEM medium, pH 7.4, with 5 mM HEPES, supplemented with 5 mM glucose, 5 mM pyruvate and 4 mM L-glutamine. Oxygen consumption and extracellular acidification rates were measured by Seahorse XF analyser and normalised using a cell viability assay, CyQUANT^™^ NF (ThermoFisher, C35006).

### Proliferation and viability assays

Proliferation of cells in real time was monitored using the xCELLigence RTCA SP instrument (Agilent). 2500-10,000 cells per well were seeded at the start of the assay, and cellular proliferation and viability was recorded automatically through increases in electrical impedance over a time course up to 7 days. Alternatively, cell viability was measured manually at discrete time points by Cyquant NF (ThermoFisher, C35006). Fluorescence readings were used to approximate the relative number of viable cells in each well. ATP:ADP ratio assays were performed using an ADP/ATP ratio assay kit (Merck, MAK135).

### RNA immunoprecipitation and quantitative PCR (RIP-qPCR)

Ribonucleoprotein complexes were immunoprecipitated from cell lysates with 4.5 μg LARP1 polyclonal antibody (Abcam, ab86359) or IgG isotype control (Abcam, ab172730) following the method described by Keene *et al*.^33^ RNA was extracted with TRIzol (ThermoFisher, 15596026). Successful immunoprecipitation of LARP1 protein was confirmed by western blot. Levels of immunoprecipitated RNA obtained from four independent replicates were quantified by RT-PCR. Relative enrichment of individual mRNAs in LARP1-immunoprecipitated samples compared with IgG isotype control samples were calculated firstly by normalising against internal levels of 18S rRNA, and then plotting enrichment levels of individual mRNAs relative to that of *ACTB* mRNA.

### Electrophoretic mobility shift assay (EMSA)

ENO1-3’UTR and G6PD-CDS IDT PAGE purified oligos **(Supplementary Table S6)** containing T7 promoter were *in vitro* transcribed using HiScribe™ T7 High Yield RNA Synthesis Kit (NEB E2040). RNA was purified using TRIzol (ThermoFisher,15596026) with the DNAse I treatment step included. EMSA was performed using EMSA SYBR Green kit (Molecular Probes, E33075), specifically, 100 ng of purified RNA was incubated with the indicated amounts of purified LARP1 protein in binding buffer supplemented with 10% [v/v] glycerol and 0.1 mg/mL BSA. The reaction mixture was separated on an 8% native polyacrylamide gel and visualized using a ChemiDoc Imaging system (BioRad) with SYBR Green detection filter.

### Immunofluorescence microscopy

HEY-A8 or OVCAR-8 cells were seeded on glass coverslips in six well plates at (2 × 10^5^ cells per well), and were transfected with siRNAs as previously described. 24 h following transfection, cells were subjected to either serum starvation or 1 μM rapamycin for 24 h followed by fixation and permeabilization with ice-cold methanol (15 min). After 3 washes with PBS, cells were blocked with 3% goat serum in PBS, followed by 1h incubation in primary antibodies LAMP1 (Abcam, ab25630), mTOR (Cell Signalling Technologies CST2983) (both 1:100 in blocking buffer). After washing in PBS, cells were incubated for 1 h in goat anti-mouse Alexa-Fluor 488, goat anti-rabbit Alexa Fluor 647 (Life Technologies A11001 and A21244) and nuclear stain DAPI at 1:400 dilution in blocking buffer. Cells were mounted on slides using ProLong Gold Anti-fade reagent (Life Technologies P36930),and visualized using 710 Confocal Microscope using 63x magnification.

### Nascent protein detection by CLICK-iT and streptavidin immunoprecipitation

Labelling was performed using the CLICK-iT labelling system (ThermoFisher, C10276) according to the manufacturer’s protocol. In brief, 24 h following transfection with siRNAs targeting LARP1 or scrambled siRNAs, cells were starved of L-methionine and treated with L-Azidohomoalanine (AHA) (ThermoFisher, C10102) for 6 h to label nascent proteins. L-AHA labelled proteins were biotinylated using PEG4 carboxamide-Propargyl Biotin (ThermoFisher, B10185) and biotin-labelled proteins were isolated by immunoprecipitation using streptavidin magnetic beads (NEB, S1420S) in an immunoprecipitation buffer containing 25 mM Tris-HCl, pH 7.6, 1 M NaCl, 1% NP40, 1% sodium deoxycholate, 0.5% SDS. Proteins were eluted in 1x Bolt LDS sample buffer (ThermoFisher, B0007) supplemented with 1x Bolt reducing agent (ThermoFisher, B0009). Eluted nascent proteins and input fractions were detected by western blot.

### Rapid method for the immunopurification of lysosomes (LysoIP) from HA-TMEM192 HeLa cells

The pLJC5-TMEM192-3xHA construct used to produce cell lines stably expressing TMEM192-3xHA was a gift from David Sabatini (Addgene plasmid # 102930; http://n2t.net/addgene:102930; RRID:Addgene_102930).

Lentiviruses were produced by transfecting HEK-293T in combination with pAmphoRand CMV-Δ8.2R packaging plasmids. Fifty hours after transfection, the virus containing supernatant was collected, centrifuged at 1,000 x g to remove cells and filter with 0.45um PVDF filters. To establish stably expressing cell lines, 500,000 HeLa cells were plated in 6-well plates in DMEM with 10% FBS and 8 μg/mL polybrene and infected with 250 μL of virus-containing media. Sixteen hours later, the media was refreshed and puromycin was added for selection. Stable cell lines were tested for the proper localization of the lysosomal tag using immunofluorescence.

8 million cells in 10 cm plates were used for each LysoIP. Cells were quickly rinsed twice with PBS and then scraped in one mL of KPBS (136 mM KCl, 10 mM KH2PO4, pH 7.25 was adjusted with KOH, as described and centrifuged at 500 x g for 3 min at 4°C.^34^ Pelleted cells were resuspended in 950 μL, and 25 μL (equivalent to 2.5% of the total cells) was reserved for further processing of the whole-cell fraction. The remaining cells were gently homogenized with 20 strokes of a 2 ml homogenizer. The homogenate was then centrifuged at 1000 x g for 2 min at 4°C and the supernatant containing the cellular organelles including lysosomes was incubated with 25 μL of KPBS prewashed anti-HA magnetic beads on a gentle rotator shaker for 15 min. Immunoprecipitates were then gently washed four times with KPBS on a DynaMag Spin Magnet. Elution was done in 30ul of 2× Laemmlli buffer at 95°C for 10 min with intense shaking and regular vortexing.

### Drug inhibition assays

Wild-type or LARP1 knockout OVCAR-8 cells were treated with rapamycin (Cambridge Bioscience, SM83-5), Torin-1 (Stratech, A8312-APE) or BEZ235 (Cambridge Bioscience, B1996). Cellular viability was measured 72 h after drug addition with CyQUANT NF. Relative IC_50_ values calculated from a log([drug]) versus normalized response curve fit using GraphPad Prism version 9 (GraphPad Software).

### Immunohistochemical staining of human tissue

An ovarian carcinoma tissue microarray was stained for LARP1 (Abcam, ab86359) and ENO1 (Abcam, ab155102). Automated staining was carried out with the Leica BOND-MAX autostainer (Leica, Microsystem) with antigen retrieval at 100ºC for 20 min with Epitope Retrieval Solution 1 or 2 respectively (AR9961, AR9640, Leica Biosystems). TMAs were blind-scored by multiplying staining intensity with staining coverage. Non-cancerous regions of cores were ignored for scoring purposes.

Multiplex (MP) immunofluorescence (IF) staining was carried out on 4 μm thick formalin fixed paraffin embedded (FFPE) sections of ovarian carcinoma or normal fallopian tube sections using the OPAL™ protocol (AKOYA Biosciences) on the Leica BOND RX^m^ autostainer (Leica, Microsystems). Staining cycles were performed using LARP-1, Abcam (ab86359)-Opal 540 and pan-cytokeratin, DAKO (M3515)-Opal 620. Primary antibodies were incubated for 30 minutes and detected using the BOND™ Polymer Refine Detection System (DS9800, Leica Biosystems) as per manufacturer’s instructions, substituting the DAB for the Opal fluorophores, with a 10-minute incubation time. Antigen retrieval at 100° C for 20 min, as per standard Leica protocol, with Epitope Retrieval (ER) Solution 2 (AR9640, Leica Biosystems) was performed before each primary antibody was applied. For LARP-1, ER Solution 1 (AR9961, Leica Biosystems) was used. Sections were then incubated for 10 minutes with spectral DAPI (FP1490, Akoya Biosciences) and the slides mounted with VECTASHIELD® Vibrance™ Antifade Mounting Medium (H-1700-10, Vector Laboratories). Whole slide scans and multispectral images (MSI) were obtained on the AKOYA Biosciences Vectra® Polaris™. Batch analysis of the MSIs from each case was performed with the inForm 2.4.8 software provided. Batched analysed MSIs were fused in HALO (Indica Labs), to produce a spectrally unmixed reconstructed whole tissue image. Following multiplex imaging, coverslips were removed and automated staining for P53 was carried out with the Leica BOND-MAX autostainer (Leica, Microsystem) using the following conditions: antigen retrieval at 100ºC for 20 min with Epitope Retrieval Solution 2 (AR9640, Leica Biosystems); primary antibody incubation with anti-P53, clone DO-7 (M7001, DAKO) and the BOND™ Polymer Refine Detection System (DS9800, Leica Biosystems) as per manufacturer’s instructions. Finally, slides were stained with the haematoxylin and eosin stain kit (H-3502, Vector).

## Results

### LARP1 regulates enzymes required for ATP generation and macromolecule synthesis

To investigate the role of LARP1 in the regulation of global protein synthesis of cancer cells, we conducted a comprehensive quantitative proteomic mass spectrometry analysis following LARP1 depletion. A total of 4621 proteins were identified, of which 907 were significantly upregulated and 1021 significantly downregulated (**Supplementary Table S1**). On clustering all identified proteins by function, we found that metabolism-related proteins displayed a strong bias towards downregulation, specifically those involved in the pentose phosphate pathway, glycolysis, gluconeogenesis, Krebs cycle, and mitochondrial-driven aerobic respiration (**Figure 1a, Supplementary Figure S1a)**.

**Figure 1.**
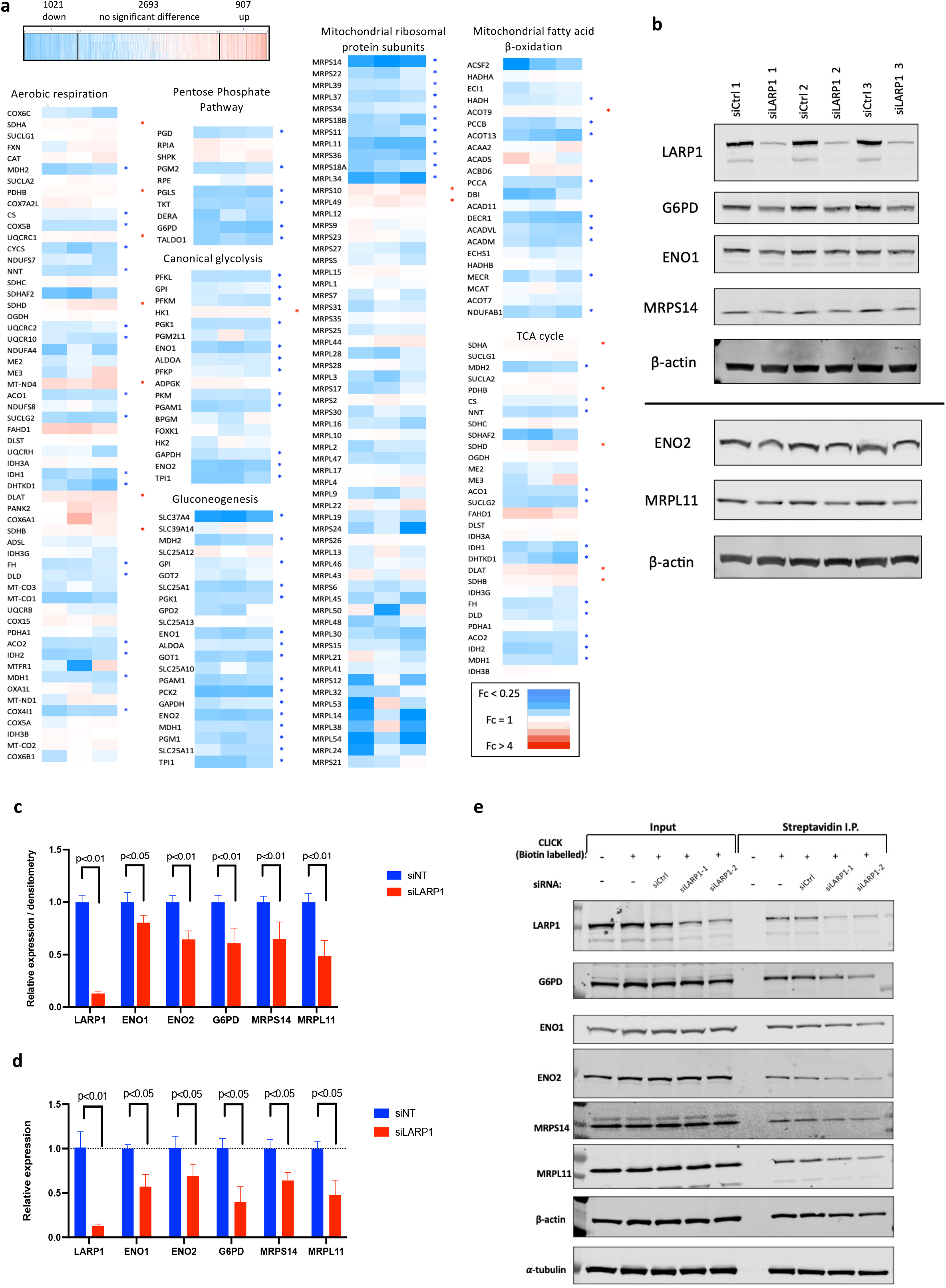
LARP1 depletion causes widespread downregulation of metabolic enzymes and mitochondrial proteins. (a) LC-MS proteomic analysis of OVCAR-8 cells following LARP1 knockdown (n=3 for control and knockdown groups). 4621 identified proteins were classified on whether protein level decreased (blue) or increased (red) in each LARP1 knockdown replicate sample compared to the control mean. 1021 proteins were identified as significantly decreased (q-value <0.05), 907 significantly increased following LARP1 knockdown (q-value <0.05). Proteins were clustered by Gene Ontology-Biological process, or gene ontology-cellular compartment with selected metabolic processes shown. Relative expression of individual proteins in each of the three LARP1 knockdown replicates compared to the mean of control samples is indicated by colour scale. Proteins showing significant increases are indicated in red, significant decreases with blue. (b) OVCAR-8 lysates after LARP1 knockdown with three different siRNA were analysed by western blot for levels of proteins identified in the proteomics screen.β-actin was used as a loading control. One experiment of three shown. (c) Densitometry analysis of blots across three different experiments with three different siRNAs targeting LARP1 showing the mean protein expression level of control samples and LARP1 knockdown samples. (d) Levels of mRNA encoding proteins identified as differentially regulated following LARP1 knockdown were analysed by quantitative RT-PCR. *ACTB* mRNA was used as a normalisation control. (e) Levels of nascent protein production were measured by CLICK-chemistry biotin labelling, followed by streptavidin pulldown of biotinylated proteins and detection by western blot. Cells were labelled with L-Azidohomoalanine (AHA) for a 6 h period between 24 h and 30 h after siRNA transfection. After biotin-labelling of L-AHA-containing proteins, these proteins were immunoprecipitated with streptavidin beads and quantified by western blot. Shown is a representative blot from one of three independent experiments.

To validate these findings, we selected a target validation set consisting of 5 proteins that were identified as significantly downregulated following LARP1 depletion in our proteomics dataset. These comprised a mixture of metabolic enzymes and mitochondrial proteins: Enolase 1 (ENO1) and Enolase 2 (ENO2) which catalyse the conversion of 2-phosphoglycerate to phosphoenolpyruvate, the critical pentose phosphate pathway enzyme glucose-6-phosphate dehydrogenase (G6PD) and the mitochondrial ribosomal subunit proteins MRPS14 and MRPL11. We then depleted LARP1 in OVCAR-8 cells across three independent assays, each using three different siRNAs targeting LARP1 and measured expression levels of these proteins (**Figure 1b**). Consistent with the proteomics data, LARP1 knockdown resulted in small but consistent decreases in all five proteins with significance confirmed by western blot densitometry normalised to β-actin (**Figure 1c**). Consistent with the role of LARP1 in regulating stability of its mRNA targets, levels of *ENO1, ENO2, G6PD, MRPS14 and MRPL11* mRNA were also reduced following LARP1 knockdown (**Figure 1d**).

Due to the relatively small changes in total levels of these metabolic and mitochondrial protein targets, we measured the level of nascent protein production following LARP1 depletion. Using CLICK chemistry to label nascent proteins with biotin, followed by streptavidin immunoprecipitation of labelled proteins, we were able to quantify protein production during a 6 h window between 24 −30 h following transfection with LARP1 siRNAs. We were thus able to distinguish protein production exclusively after LARP1 was depleted from overall protein levels. During this 6 h time window, the production of nascent LARP1, G6PD, ENO1, ENO2, MRPS14 and MRPL11 was substantially reduced in LARP1-depleted samples compared with control samples (**Figure 1e**). By contrast production of nascent β-actin and α-tubulin production was not significantly different in control samples versus LARP1-depleted samples. Collectively, these results indicate that following LARP1 depletion, while global protein synthesis is not downregulated, synthesis of metabolic and mitochondrial proteins is selectively reduced.

As an RNA-binding protein, we hypothesised that LARP1 exerted control over expression of these proteins by binding and regulating the stability and translation of their encoding mRNAs. In addition to binding TOP mRNAs, multiple studies have demonstrated that LARP1 binds thousands of non-TOP mRNAs, and that these non-TOP mRNAs comprise between 94-98% of the total mRNA interactome of LARP1.^10,29,35,36^ To determine if mRNAs bound by LARP1 showed enrichment for particular biological processes that might drive cancer progression, including metabolism, we analysed a previously described list of 2658 LARP1-bound mRNAs, coding for known proteins (**Supplementary Table S2**).^29^ Genes were stratified by gene ontology and biological process using the PANTHER gene classification tool (http://www.pantherdb.org). Out of 89 statistically significant enriched pathways, 26 were linked to catabolic or metabolic processes with a further 24 to biosynthetic processes (**Figure 2a, Supplementary Table S3**), demonstrating the importance of LARP1 in binding and regulating metabolism-associated mRNAs.

**Figure 2.**
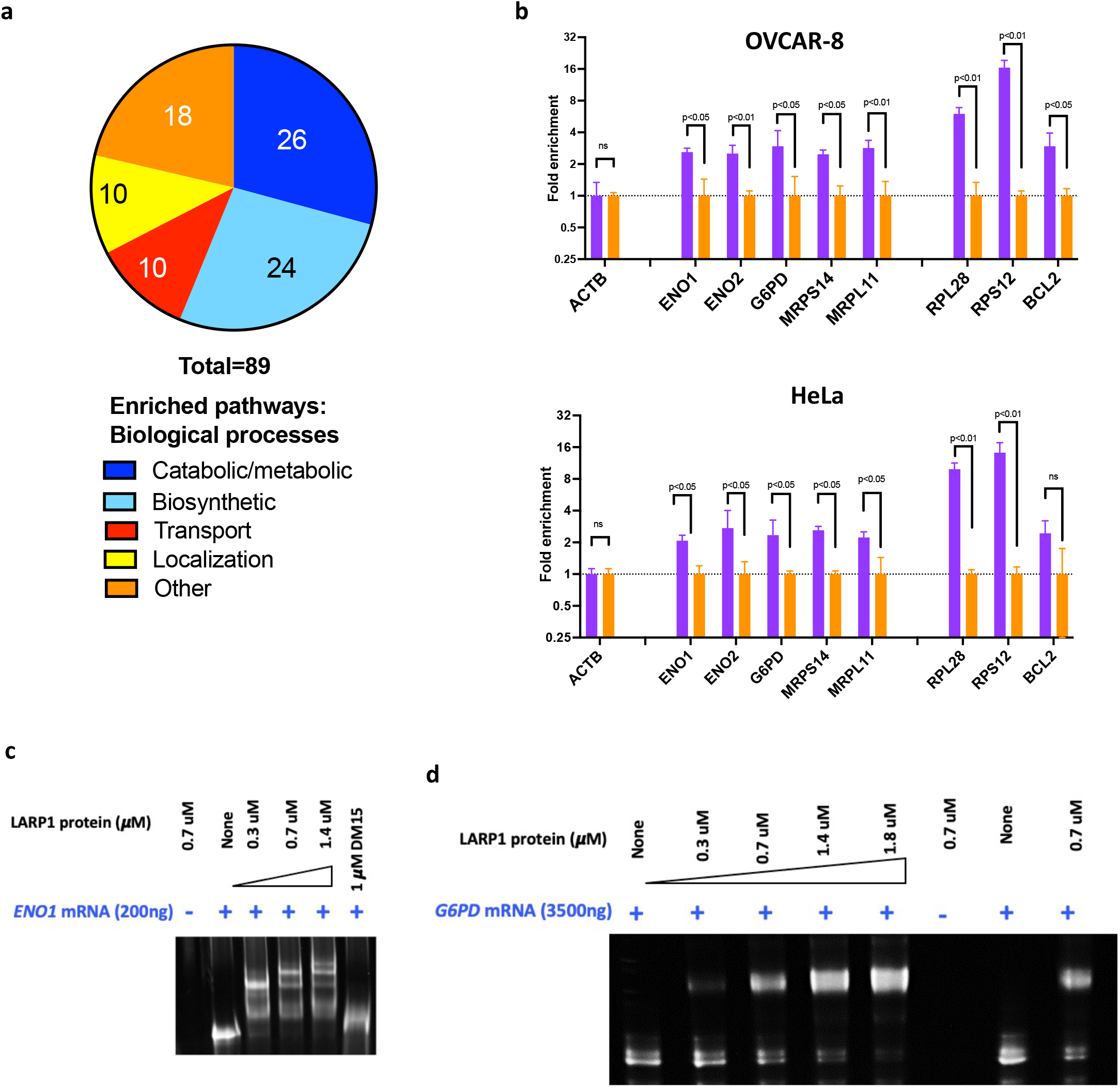
LARP1 directly binds mRNAs encoding metabolic enzymes. (a) Biological processes identified as enriched in LARP1-bound mRNAs (in our previously reported LARP1 interactome, Supplementary Table S2) were stratified by class. (b) RNA immunoprecipation followed by qRT-PCR was used to quantify the level of binding of individual mRNAs to LARP1 in OVCAR-8 and HeLa cells. Quantification of the level of enrichment of individual mRNAs bound to LARP1 was calculated relative to *18S* rRNA levels immunoprecipitated by IgG isotype control antibodies and normalized against relative enrichment of *ACTB* mRNA. Enriched LARP1-binding of an mRNA compared to *ACTB* is indicated by Fc>1, with significant binding requiring Fc>2. (c) Electrophoretic mobility shift assay (EMSA) on region corresponding to the 3’ UTR of *ENO1* mRNA showing shift in binding when incubated with increasing concentrations of LARP1. Incubation with the DM15 region of LARP1 alone did not induce a shift in *ENO1* mRNA migration. (d) EMSA demonstrating a shift in migration of a region of the CDS of *G6PD* mRNA with increasing concentrations of LARP1. Pairwise comparisons performed using Kolmogorov-Smirnov test *, (p<0.05), ** (p<0.01)

To confirm whether LARP1 directly binds the mRNAs encoding the proteins whose expression reduced after LARP1 depletion, we performed an RNA-immunoprecipitation assay in OVCAR-8 and HeLa cells using an anti-LARP1 antibody to immunoprecipitate ribonucleoprotein complexes, (**Supplementary Figure 1b**). Enrichment of individual mRNAs was calculated relative to *ACTB* mRNA, which does not show enriched binding to LARP1.^29^. The selected metabolic genes *ENO1, ENO2, G6PD, MRPS14* and *MRPL11* were all enriched following immunoprecipitation of LARP1 in both OVCAR-8 and HeLa cells, confirming binding by LARP1 (**Figure 2b**). Consistent with previously published findings, we also demonstrated that mRNAs encoding the ribosomal proteins RPL28 and RPS12, as well as the apoptosis-regulating BCL2, were enriched for binding to LARP1.^10,29,35,36^

We selected two targets displaying enriched binding (*ENO1* and G6PD) for validation by electrophoretic mobility shift assays (EMSA). By incubating *in vitro* transcribed 3’-UTR regions of *ENO1* mRNA with LARP1 protein, we confirmed direct binding, with greater shifts in ENO1 binding at higher concentrations of LARP1 (**Figure 2c**).^27^ We also showed in this EMSA that the DM15 domain of LARP1 alone is not sufficient for *ENO1* binding, as this failed to cause a shift in migration. To determine if *G6PD* mRNA bound LARP1, we used using a CDS region of *G6PD*, previously identified as binding LARP1in a large LARP1 iCLIP dataset.^35^ We demonstrated by EMSA that increasing levels of LARP1 resulted in a gradually increasing shift of *G6PD* mRNA (**Figure 2d**). As with *ENO1*, only full-length LARP1, and not the DM15 domain alone, was capable of binding *G6PD* (**Supplementary Figure 1c**). While the binding of TOP mRNAs depends solely on DM15,^15^ this demonstrates that non-TOP mRNAs such as *G6PD* and *ENO1* are dependent on other domains of LARP1, such as the La module, for binding.

### LARP1 drives high-level metabolic activity in cancer cells

To assess the impact of LARP1 depletion on the global metabolome, we conducted a metabolomics screen on OVCAR-8 cells following LARP1 knockdown. A principal components analysis confirmed that LARP1 knockdown significantly altered the global metabolisms of the cells (**Figure 3a**), while a hierarchically-clustered abundance heatmap shows distinct patterns of clustered metabolites abundance changes between the two experimental groups (**Figure 3b**). Our analysis of individual pathways revealed significant alterations in metabolite levels across the glycolytic, Krebs cycle and pentose phosphate pathways (**Figure 3c**). Primarily, this included altered metabolite abundance in pathways related to glucose and glutamine catabolism for ATP generation and also a relative depletion of lactate, indicating reduced glycolysis following LARP1 knockdown. LARP1 knockdown also resulted in alterations in glutamine and glutamate metabolism (**Figure 3d)**, collectively implying impaired ability to generate ATP or to perform anabolism. The observed disruption to the pentose phosphate pathway and accumulation of ribose-5-phosphate indicated defective processing of nucleotides for DNA and RNA production.

**Figure 3.**
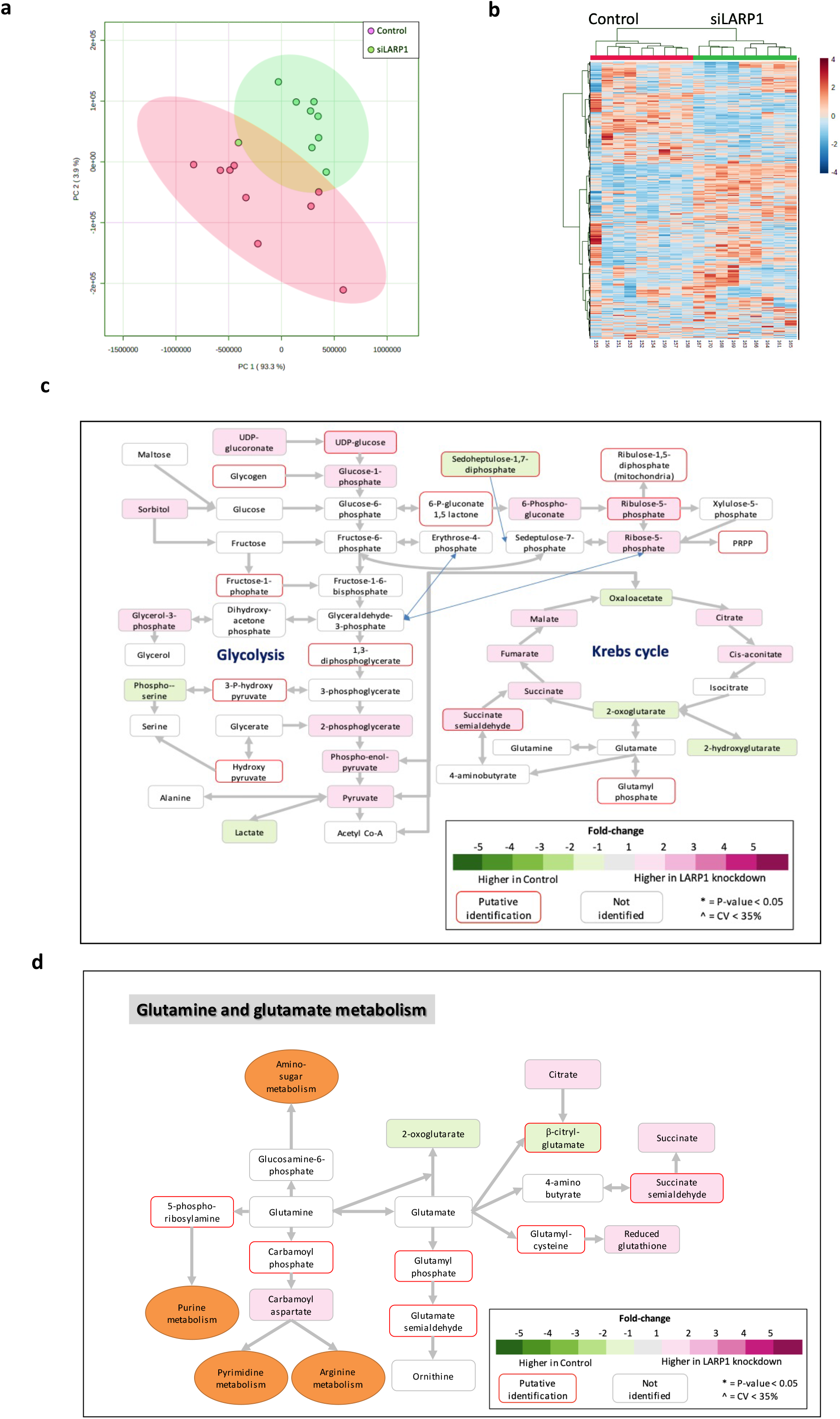
The global metabolome of cancer cells is altered following LARP1 depletion. Metabolite levels in control or LARP1-knockdown OVCAR-8 cells were measured by mass spectrometry in a metabolomics screen (n=9). (a) Principal components analysis of individual replicates from the control and LARP1 knockdown groups showing significant differences in the metabolomic profile following LARP1 knockdown. (b) Hierarchically-clustered abundance heatmap of metabolites in control and LARP1 knockdown groups clustered by pathways (c,d) Changes in metabolite levels following LARP1 knockdown across metabolic networks and are indicated for the Krebs cycle, glycolytic and pentose phosphate pathways (c) and for glutamine/glutamate metabolism (d). Increased levels following LARP1 knockdown are marked in pink, decreased levels are marked in green.

Consistent with this, we observed widespread changes in purine and pyrimidine metabolism (**Supplementary Figure S2a**,**b**), suggesting an effect on nucleic acid synthesis.

To investigate the biological importance of LARP1 in metabolic regulation of cancer cells, we depleted LARP1 in four cancer cell lines (the ovarian carcinoma lines OVCAR-8 and SKOV3, the osteosarcoma line U2OS and the cervical carcinoma line HeLa) and performed Seahorse MitoStress assays to measure aerobic respiration and glycolysis (**Figure 4a**). Across all four cell lines, a significant reduction in basal oxygen consumption rate (OCR) was observed on LARP1 knockdown (**Figure 4b and Supplementary Figure S3a**). Extracellular acidification rates (ECAR) (approximating glycolysis) were also significantly reduced following LARP1 knockdown in OVCAR-8, SKOV3 and HeLa cells (**Figure 4b**), and thus increased glycolysis was not able to compensate for the observed reduction in aerobic respiration. Consistent with these data, we observed reduced proliferation rates following LARP1 knockdown in all four cell lines (**Supplementary Figure S3b**).

**Figure 4.**
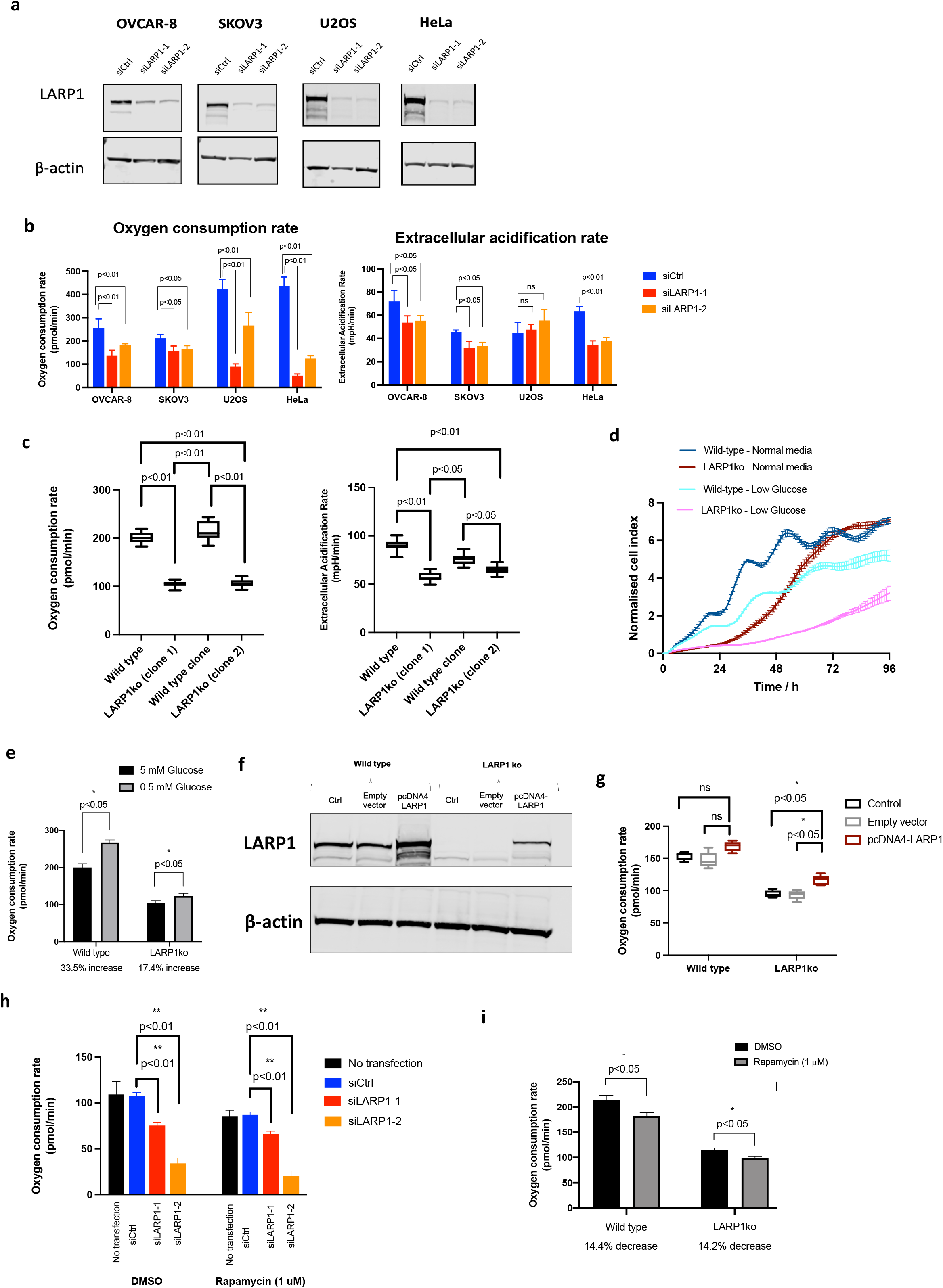
LARP1 depletion causes reduced metabolic activity of cancer cells independent of mTORC1 status. (a) LARP1 was depleted across four different cell lines using two different siRNAs confirmed by western blot after 48 h. (b) Basal oxygen consumption rates (OCR) and basal extracellular acidification rate (ECAR) of LARP1-depleted cells were measured by Seahorse Mitostress assays. Full assay results (following sequential addition of oligomycin, FCCP and antimycin A/rotenone) are shown in Supplementary Figure S2a. (c) Basal OCR and ECAR rates for LARP1-knockout OVCAR-8 compared with wild type cells (d) Proliferation rates of wild type and LARP1ko OVCAR-8 cells grown in normal (5 mM) or low glucose (0.5 mM) measured using Xcelligence real time cell analysis. (e) Basal OCR of wild-type versus LARP1ko cells when grown in 5 mM versus 0.5 mM glucose. (**f**,**g**) A LARP1 expression construct (pcDNA4-LARP1) was transfected into LARP1ko OVCAR-8 cells to determine if the metabolic phenotype could be rescued. (f) Successful expression from the construct was confirmed by western blot. (g) Basal oxygen consumption rates of wild-type and LARP1ko OVCAR-8 cells transfected with the LARP1 expression construct or an empty vector. The LARP1 expression vector was able to achieve a partial restoration of the OCR induced by LARP1 knockdown. Pairwise comparisons performed using Kolmogorov-Smirnov test *, (p<0.05), ** (p<0.01) (**h)** OCR was measured by Seahorse Mitostress assay OVCAR-8 cells were pre-treated with 1 μM rapamycin 24 h prior to transfection with LARP1 siRNAs and then for a further 48 h. (i) Oxygen consumption rates of wild type or LARP1-ko OVCAR-8 cells were measured with or without 1 μM rapamycin (48 h).

To confirm that LARP1 knockout by CRISPR-Cas9 induced a similar phenotype to siRNA knockdown, we compared the metabolism of LARP1-knockout clones to wild-type OVCAR-8 cells. Through monoclonal selection with puromycin, we thus generated two clones, expressing no full length LARP1 (albeit with low-level expression of the minor isoform of LARP1),^37^ along with an additional clone which we confirmed to be genetically identical to wild-type, which we used an additional control for any phenotypic changes induced during the clonal selection process. (**Supplementary Figure 3c**).

Both LARP1 knockout clones exhibited significantly reduced basal oxygen consumption and extracellular acidification rate compared with wild type, indicating reduced aerobic respiration and glycolysis respectively **(Figure 4c)**. We have demonstrated previously that LARP1 depletion results in both slower proliferation and higher levels of apoptosis in cancer cells.^27,29^ To determine if this phenotype was amplified by nutrient stress, we compared the effects of glucose deprivation in wild-type and LARP1ko (clone 1) OVCAR-8 lines. As expected, glucose deprivation (5 mM to 0.5 mM) resulted in reduced proliferation rates, with a disproportionately greater effect on LARP1ko cells than wild-type (**Figure 4d**), indicating greater adaptability of wild-type cells to low glucose conditions. We then questioned whether the failure to adapt to low glucose was due to a failure to increase aerobic respiration to compensate for reduced glycolysis. Wild type OVCAR-8 cells were able to significantly increase aerobic respiration rates (by 33.5%) when glucose concentrations were reduced to 0.5 mM, but LARP1ko cells only displayed a significantly smaller increase of 17.4% (p<0.05) (**Figure 4e**). A similar disparity in response to glucose deprivation (25.7% versus 12.1% increase was observed in OVCAR-8 cells where LARP1 was depleted by siRNA-mediated knockdown (**Supplementary Figure S3d)**. This indicates that LARP1 contributes to the metabolic plasticity of cancer cells, allowing them to adapt to nutrient-poor environments.

To confirm that the reduced oxidative metabolism in LARP1ko cells was directly attributable to loss of LARP1, we performed rescue with a LARP1 expression vector transfected into LARP1ko OVCAR-8 cells (**Figure 4f**). Transfection with the LARP1 expression vector did not significantly alter OCR in wild-type cells but did result in a significant increase in OCR in LARP1ko cells (**Figure 4g**). A similar assay, combining LARP1 expression vector transfection with siRNA mediated knockdown rather than CRISPR-Cas9 stable knockout, also resulted in a partial rescue of oxygen consumption rates (**Supplementary Figure S3e**,**f**). Due to the observed reduction in oxygen consumption rates on LARP1 knockdown, we hypothesised that mitochondrial morphology may have been affected. However, electron microscopy across multiple OVCAR-8 cell sections showed no visible defects in mitochondria following LARP1 knockdown (**Supplementary figure S4**). Any effect on mitochondrial respiration was therefore the result of impaired mitochondrial function, rather than changes in size or morphology.

### LARP1 promotes metabolism in cancer cells independent of mTORC1 activation status

The best characterised role of LARP1 is as a downstream phospho-target of mTORC1 in regulating the initiation of translation of TOP mRNAs ^12,38^. Therefore, we next addressed whether the observed effect of LARP1 depletion on metabolism and proliferation was dependent on mTORC1 being active. Rapamycin and mTORC1 inhibition are known to alter the phosphorylation status of at least 26 LARP1 residues, resulting in release of TOP mRNA binding when mTORC1 is active.^16^ However, the role of mTOR in control of non-TOP mRNA binding is less clear, with some constitutively bound to LARP1 and others whose binding is mTOR-dependent.^36^ To address whether the effect of LARP1 on metabolic activity relied on mTORC1 being in an active state, we treated OVCAR-8 cells with rapamycin prior to and following siRNA-mediated LARP1 depletion. We confirmed that rapamycin had successfully induced mTORC1 inhibition through ribosomal protein S6 dephosphorylation (**Supplementary Figure S5a)**. LARP1 knockdown with two different siRNAs induced a significant reduction in oxygen consumption rates indicating but this effect was observed whether or not cells were treated with rapamycin prior to and during the assay (**Figure 4h**). While rapamycin caused only a slight reduction in proliferation rate, the two different LARP1 siRNAs substantially reduced cellular proliferation whether or not rapamycin was present (**Supplementary Figure 5b**). This suggested that mTORC1 activity was not required in order for the effect of LARP1 depletion on cancer cell metabolism and viability.

Having determined that the role of LARP1 in metabolic regulation is mTORC1-independent, we addressed the question in reverse – whether LARP1 is required for cells to respond to changes in mTORC1 activity, and whether LARP1ko are equally sensitive to mTORC1 inhibitors. As LARP1 binding to TOP mRNAs is regulated by mTORC1, we speculated that LARP1ko cells may be less sensitive to rapamycin than wild-type cells. We therefore treated OVCAR-8 LARP1ko cells with rapamycin for 48-120 h, combined with assays to measure oxygen consumption and proliferation rates. Although, rapamycin treatment (1 μM) caused a significant reduction in the proliferation rates of both wild-type and LARP1ko OVCAR-8 cells over a 120 h time course, wild type cells showed no greater reduction than LARP1ko cells (**Supplementary Figure S5c**). Similarly, while rapamycin induced a significant decrease in the oxygen consumption rate of both wild-type and LARP1 knockout cells, this decrease was not significantly different between wild-type and LARP1ko (14.4% and 14.2% respectively) (**Figure 4i**). To confirm, that no difference in sensitivity to mTORC1 inhibition existed between wild-type and LARP1-depleted cells, we measured the IC_50_ of rapamycin and the mTORC1/2 inhibitors Torin-1 and BEZ235. None of the three displayed significant differences in the inhibition curves or IC_50_ values between wild-type and LARP1 knockout cells (**Supplementary Figure S5d-f**). This indicates that the effect of mTOR inhibitors on oxidative respiration and cellular proliferation is not affected by the presence or absence of LARP1. Collectively, these data demonstrate that the role of LARP1 in regulating metabolism in cancer cells is therefore independent of their mTORC1 activity status.

### LARP1 promotes mTORC1 activation by regulating the localisation of mTORC1 on the lysosomal surface

The process by which mTORC1 is trafficked to the lysosomal surface is controlled by the interplay between amino-acids and the highly ATP sensitive vacuolar-associated enzyme v-ATPase.^7^ To explore whether mTORC1 localisation was altered by LARP1, we analysed the localisation of mTOR in the ovarian cancer cell lines HEY-A8 and OVCAR-8 cells. Using LAMP1 as a lysosomal marker, we showed that mTOR colocalises with the lysosome when LARP1 is present but that this localisation was reduced when LARP1 was depleted by siRNA (**Figure 5a**). Furthermore, the effect of LARP1 depletion is still observed when mTOR is inhibited either by serum starvation or 1 μM rapamycin treatment. Although both serum starvation and rapamycin treatment reduced colocalization of LAMP1 with mTOR, in both cases, this was reduced still further following LARP1 knockdown.

**Figure 5.**
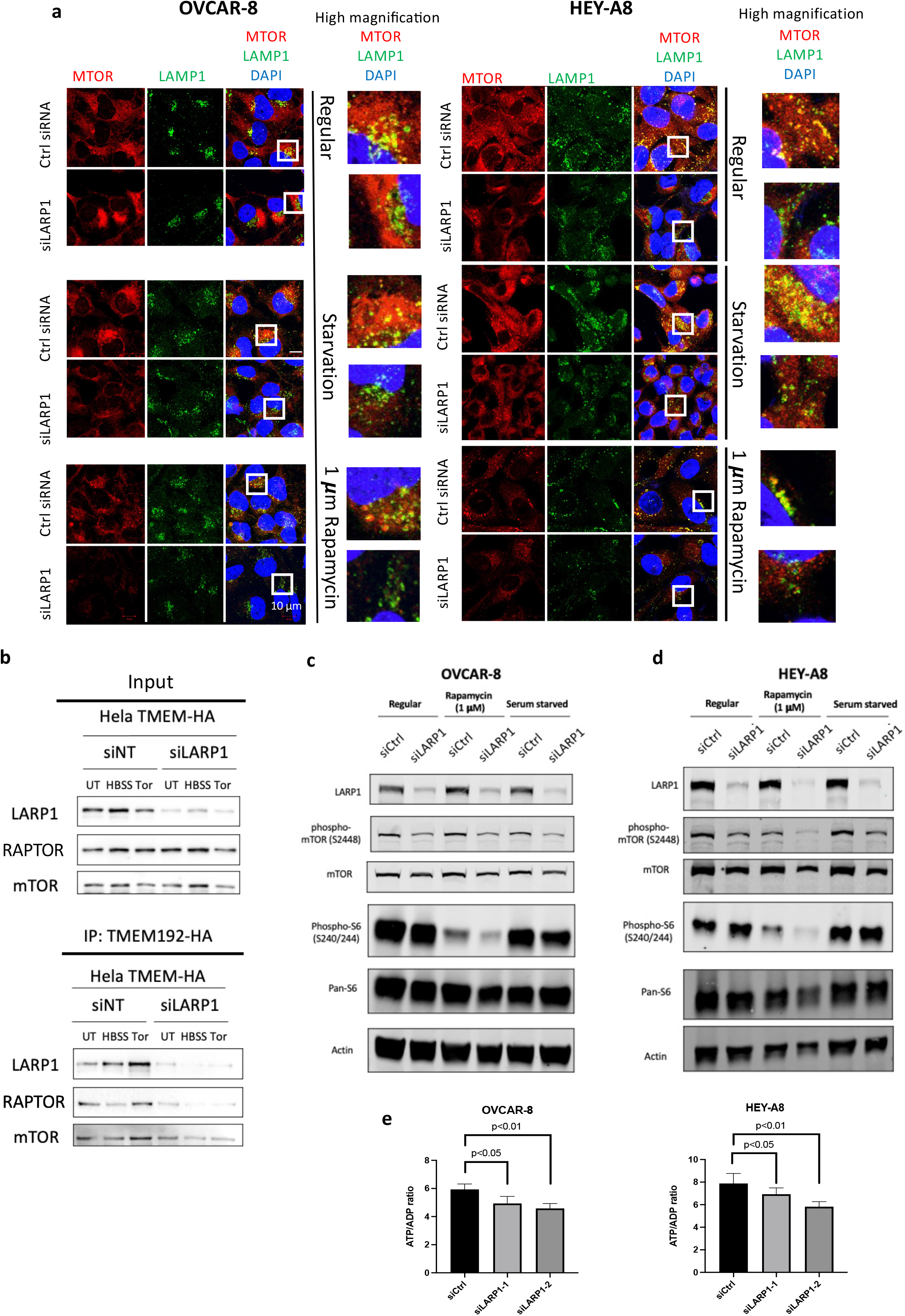
LARP1 depletion prevents mTOR localization to the lysosome and reduces mTORC1 activity. (a) mTOR (red), DAPI (blue), and the lysosomal protein LAMP1 (green) were visualised in OVCAR-8 and HEY8 cells by immunofluorescence microscopy following LARP1 depletion (48 h after siRNA transfection). Cells were grown in normal media, serum-depleted media for 24 h, or media containing 1 μM rapamycin for 24 h to inhibit mTORC1. Colocalisation of mTOR and LAMP1 is indicated by yellow staining in the merged image. (b) HeLa cells stably expressing HA-TMEM192 were treated with siRNAs targeting LARP1 for 48 h. Anti-HA immunoprecipitation was used to identify proteins bound to lysosomal TMEM192, with or without starvation by growth in Hank’s buffered saline solution (HBSS) or treatment with Torin-1. Blots shown are representative of three independent assays. (c,d) Level of mTORC1 activation in OVCAR-8 and HEY-A8 cells following LARP1 depletion by siRNA, as measured by phosphorylation of mTOR, AKT1 and ribosomal protein S6. Cells were transfected with siRNAs 48 h prior to imaging and grown in normal, serum-depleted or rapamycin (1 μM)-containing media for 24 h prior to imaging. (e) ATP:ADP ratios in OVCAR-8 and HEY-A8 cells 48 h after siRNA transfection.

To confirm that LARP1 was affecting the localisation of mTORC1 components we utilised a cell line model in which the lysosomal protein TMEM192 was stably expressed with an HA tag in HeLa cells, (*cell line generated by Pauline Lascaux in the lab of Prof. Kristjan Ramadan)*. In this model, any proteins localised at the surface of the lysosome should immunoprecipitate with HA-TMEM192. We therefore immunoprecipitated HA-TMEM192 in cells where LARP1 had been depleted by siRNA versus non-targeting siRNA (siNT) treated cells. Assays were performed either in complete media, in Hank’s buffered saline solution (HBSS) to mimic starvation conditions, or with Torin-1 treatment to inhibit mTOR activity. It has previously been shown that HBSS treatment reduces RAPTOR and mTOR binding to the lysosome while Torin-1 treatment increased lysosomal binding, and our results were consistent with this.^34^ However, across all three conditions tested (untreated, HBSS and Torin-1), RAPTOR and mTOR binding to HA-TMEM192 was reduced following LARP1 depletion., despite no substantial change in overall RAPTOR and mTOR expression levels – reflected by input levels (**Figure 5b**). This indicates that both RAPTOR and mTOR localisation at the lysosome are reduced following LARP1 depletion, and therefore that LARP1 may influence the state of mTORC1 activation.

As loss mTOR and RAPTOR localisation at the lysosome would suggest that mTORC1 activity and downstream signalling was reduced, we examined phosphorylation of downstream targets of mTORC1 in OVCAR-8 and HEY-A8 cells following LARP1 knockdown (**Figure 5c,d**). Again, different conditions were used, either rapamycin treated, serum starved or regular media. Across both cell lines, phosphorylation of mTOR at S2448 was reduced, independent of rapamycin treatment or serum starvation. In OVCAR-8 S6 phosphorylation at S240/244 was slightly reduced by LARP1 knockdown in all of regular, rapamycin-treated and serum-starved conditions. In HEY-A8 cells, we only observed an effect of LARP1 depletion on S6 S240/244 phosphorylation when combined with rapamycin treatment. Collectively, these data indicate that LARP1 positively regulates mTORC1 signalling, and that high LARP1 expression levels can potentiate mTORC1 signalling in cancer cells.

To identify possible causes of the reduced mTOR activity following LARP1 depletion, we measured the ratio of ATP to ADP in OVCAR-8 and HEY-A8 cells following LARP1 depletion **(Figure 5e)**. Consistent with previous results we demonstrated that the ATP:ADP ratio was reduced in both cell lines following LARP1 knockdown, indicating an insufficient rate of ATP production following LARP1 knockdown, and providing a possible mechanism for why LARP1 depletion results in reduced mTORC1 activation.

### LARP1 correlates with ENO1 and G6PD expression in human ovarian tumour tissue

Levels of LARP1 are frequently upregulated in ovarian cancer, and we hypothesise this drives tumour aggressiveness by promoting high-level metabolism within the tumour tissue. In ovarian tissue sections containing both regions of normal epithelium and ovarian carcinoma, levels of LARP1 were substantially higher in the tumour tissue (**Figure 6a**). To confirm association of LARP1 with its metabolic targets in tumour tissue, we analysed a human ovarian cancer tissue microarray with 894 individual tissue cores for expression levels of LARP1, ENO1 and G6PD. Cores were scored based on a combination of staining intensity and coverage within the tumour core (**Supplementary figure S6a**,**b**). Both ENO1 and G6PD displayed regional correlations in staining with LARP1, with regions higher in LARP1 also elevated in ENO1 and G6PD (**Figure 6b**). Almost no staining of either LARP1 or ENO1 was present in non-cancerous tissue within the cores. Across the entire TMA, ENO1 and G6PD staining correlated positively with LARP1 (ENO1, Spearman r=0.59, p<0.001; G6PD, Spearman r=0.44, p<0.001) (**Figure 6c,d**). Collectively, these data demonstrate the prominence of LARP1 in ovarian cancer and its correlated expression with a validated metabolic target (ENO1) *in vitro* that also exists *in vivo*.

**Figure 6.**
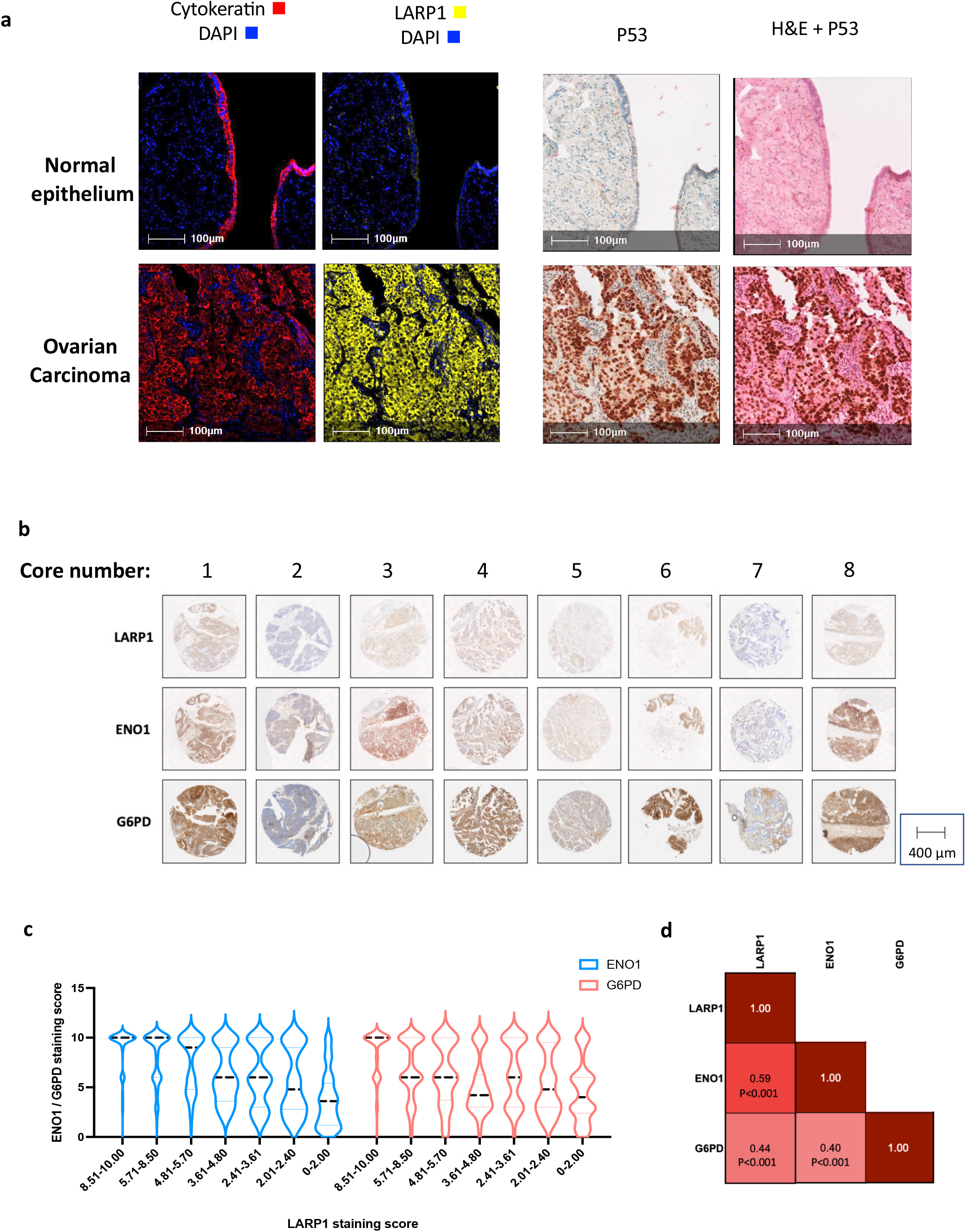
Human ovarian cancer sections express high LARP1 and are significantly correlated with expression of ENO1 and G6PD. (a) Ovarian tissue sections containing regions of normal epithelium and ovarian carcinoma were stained by multiplex immunofluorescence microscopy for LARP1 (yellow), cytokeratin (red) and DAPI (blue) (left panels), followed by P53 and haematoxylin and eosin (right panels). (b) Representative images of cores within a human tissue microarray of ovarian cancer (894 individual cores), stained with LARP1, ENO1 and G6PD. G6PD section is non-consecutive with LARP1 and ENO1. (c) Staining of LARP1, ENO1 and G6PD within the human TMA was scored for intensity and coverage, with cores assigned a score between 0 and 10. Distribution of ENO1 and G6PD scores are shown in violin plots, grouped by their LARP1 staining score. Dashed line represents the median staining score, dotted lines represent the upper and lower quartile scores. (d) Correlation matrix (Spearman) of LARP1, ENO1 and G6PD staining scores across the ovarian TMA. LARP1 was significantly positively correlated with both ENO1 and G6PD.

## Discussion

We initially set out to determine how regulation of the RNA-binding protein LARP1, downstream of mTORC1, could potentially impact metabolic signalling in cancer cells. We demonstrated that LARP1 drives high-level mitochondrial respiration and glycolysis in cancer cells through direct mRNA binding, in turn sustaining the energetic needs of cancer cells for proliferation. To our surprise we found that not only was LARP1’s regulation of metabolic signalling pathways mTORC1-independent, but more importantly that LARP1 regulates mTORC1 activation. Using LAMP1 as a lysosomal marker, we showed that mTOR colocalises with the lysosome when LARP1 is present but that this localisation is reduced when LARP1 is depleted. This shows that, through lysosomal localisation (likely as a downstream consequence of the metabolic effects of LARP1), LARP1 potentiates mTORC1 activity. LARP1 has previously been shown to be phosphorylated by mTORC1 on multiple sites to promote ribosome biogenesis, linking with our findings that LARP1 drives metabolism and increased mTORC1 activity. Our findings point towards a feed-forward loop in which LARP1 promotes ATP generation and simultaneously promotes mTORC1 signalling to drive increased ribosome biogenesis and cancer cell proliferation. In this model for the LARP1-mTORC1 axis we propose that: (1) LARP1 has an mTORC1-independent role in its regulation of metabolic factors, (2) LARP1 can drive mTORC1 activation through lysosomal localisation, (3) subsequent phosphorylation of LARP1 by activated mTORC1 drives ribosome biogenesis in cancer cells.

The anabolic demands of cancer cells for ribosome biogenesis, lipid synthesis and other related processes must be coordinated with substantial ATP generation^39,40^. In normal cells this would be tightly coupled through mTORC1 and regulatory factors, but in cancer cells this coordination is often lost, resulting in critical stress when their energetic demands are not met^4,41^. Consistent with this, we found in OVCAR-8 cells that LARP1 depletion did not result in reduced expression of ribosomal protein subunits, despite significant reductions in respiration, glycolysis and ATP:ADP ratios. Consequently, following LARP1 depletion, cancer cells displayed higher levels of stress and eventually underwent apoptosis. This supports the hypothesis that many cancer cells (such as the OVCAR-8, SKOV3, HeLa, U2OS and HEY-A8 cells used in this study) become “addicted” to LARP1 due to the metabolic boost that it drives and that the constitutively activated anabolic pathways of cancer cells must be supported by associated metabolic signalling or critical stress in the cells will result.

There are over 2,000 RNA binding proteins (RBPs) within human cells, characterised by having one or more RNA-binding domains (RBDs)^42^. An individual RBP may have interactomes of up to thousands of mRNAs, often acting within specific physiological pathways^43^. In lower organisms, RBPs are uniquely responsive to environmental stresses whereby, through modulating translation initiation by altering the conformation of bound target mRNAs, they promote adaptive survival responses such as cold shock adaptation^44^. In human cells, RBPs have housekeeping roles but their dysregulation is increasingly observed in disease states such as cancer where they act as key regulators^45^.

The RBP LARP1 is expressed at low levels in normal cells where it functions in embryogenesis, cell cycle and, importantly, in ribosomal turnover^30,46^. Following evidence that it is a direct phosphorylation target of mTORC1 and regulates ribosomal mRNA expression, LARP1 has been named the ‘missing link’ between mTORC1 activation and ribosomal biogenesis^47^. A recently proposed model is that when mTOR is inactive, LARP1 and its interacting partner poly-A binding protein (PABP) bind and repress mRNAs that encode ribosomal proteins^16^. These mRNAs carry a characteristic 42-nucleotide 5’ terminal oligopyrimidine (TOP) tract immediately adjacent to their 5’ cap^48^. By binding to this TOP motif, LARP1 prevents eIF4E from accessing the 5’ cap of these transcripts. However, when phosphorylated by mTORC1, LARP1 undergoes a conformational change that frees the 5’ cap and allows eIF4E to bind, nucleate the eIF4F complex and initiate translation^15^.

Unlike its predominantly repressive effect on TOP translation in normal cells, in cancer LARP1 has a predominantly permissive effect on the translation of its mRNA interactome. This includes pro-survival mRNAs such as BCL2 that have been shown to contribute to the apoptosis observed on LARP1 knockdown^27^. We have shown that, in addition to binding survival mRNAs, LARP1 also positively regulates the expression of multiple metabolic enzymes. This includes proteins we have validated including ENO1, ENO2, G6PD and the mitochondrial proteins MRPL11 and MRPS14 but our proteomics data also implicate LARP1 in driving the expression of hundreds more. Moreover, in cancer cells in which mTOR is hyperactive, we have shown that LARP1 regulation of these metabolic mRNAs is mTOR independent. The domain of LARP1 protein that mediates these interactions is unknown; others have shown that LARP1 binds non-TOP mRNAs using its more N-terminally located RNA-binding regions, such as the La domain and RNA recognition motif^49,50^. The site of LARP1 binding with target mRNAs also appears to differ between TOP and non-TOP mRNAs. In TOP mRNAs, LARP1 binds the 5’ pyrimidine-rich TOP sequences themselves whereas in non-TOP mRNAs, LARP1 binds other sites including 3’UTR and coding sequences^35,36,49,50^. A consensus binding sequence for LARP1 has yet to be identified, perhaps indicating that, like its family members La, LARP4a, LARP4b and LARP6, LARP1 recognises and binds structural motifs in target mRNAs rather than conserved sequences^51–54^.

Collectively, our data demonstrates that in cancer cells where mTOR is hyperactivated LARP1 acts both upstream and downstream of mTORC1. In these mTOR hyperactivated cell lines, mTORC1 is non-essential for metabolic reprogramming and that this role is assumed by the RNA binding protein LARP1. In doing so, LARP1 both drives the metabolic requirements of the cancer cell, potentiates AKT and mTORC1 signalling, drives cancer cell proliferation and prevents cell death. By driving glycolysis, pentose phosphate pathway usage and mitochondrial activity, LARP1 boosts ATP generation thereby promoting mTORC1 activation. This leads to the intriguing concept that LARP1 drives increased metabolic flux, ATP generation and mTORC1 activation as part of coordinated and overarching process of metabolic reprogramming. Thus, according to our model, LARP1 has the ability to couple ATP generation with anabolism ensuring that the proliferation of cancer cells is coupled with a means of sustaining the associated energetic requirements.

In conclusion, we have demonstrated LARP1 as a critical gate-keeper for driving the metabolic requirements and ensuring the survival and proliferation of mTORC1-driven cancer cells in hostile environmental conditions.

## Acknowledgements

We thank Ms Fenella Gross and Mrs. Arussa Nawaz (Department of Oncology, University of Oxford) for their assistance and helpful comments.

