## Supplementary Figures for "LARP1 regulates metabolism and mTORC1 activity in cancer"

### Supplementary Figure S1

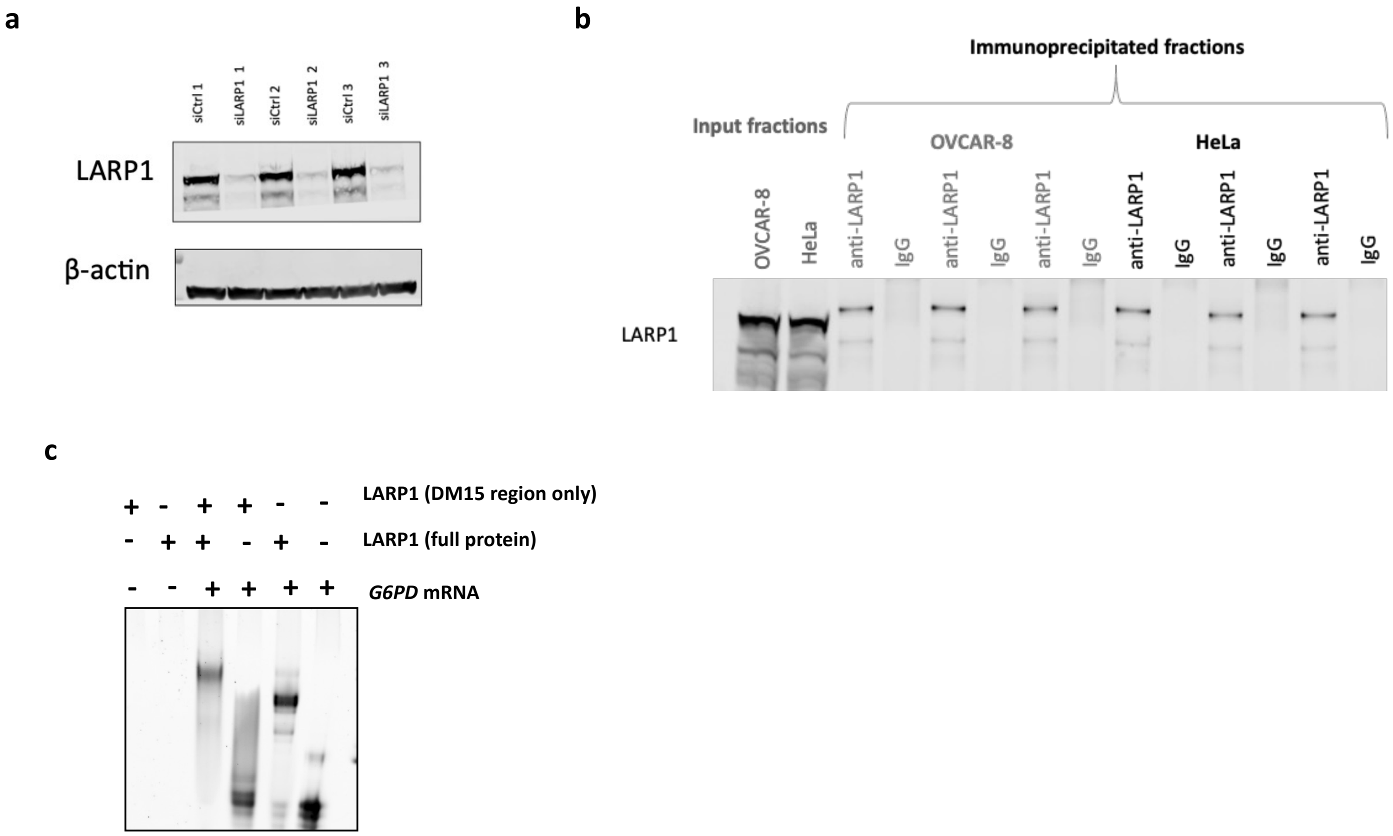

### Supplementary Figure S2

**a**

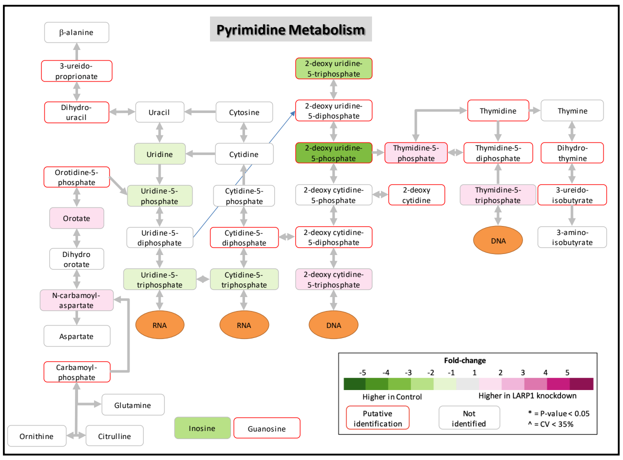


**b**

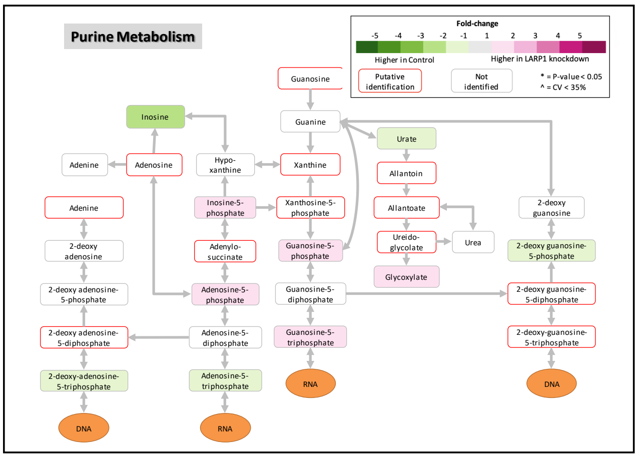

### Supplementary Figure S3

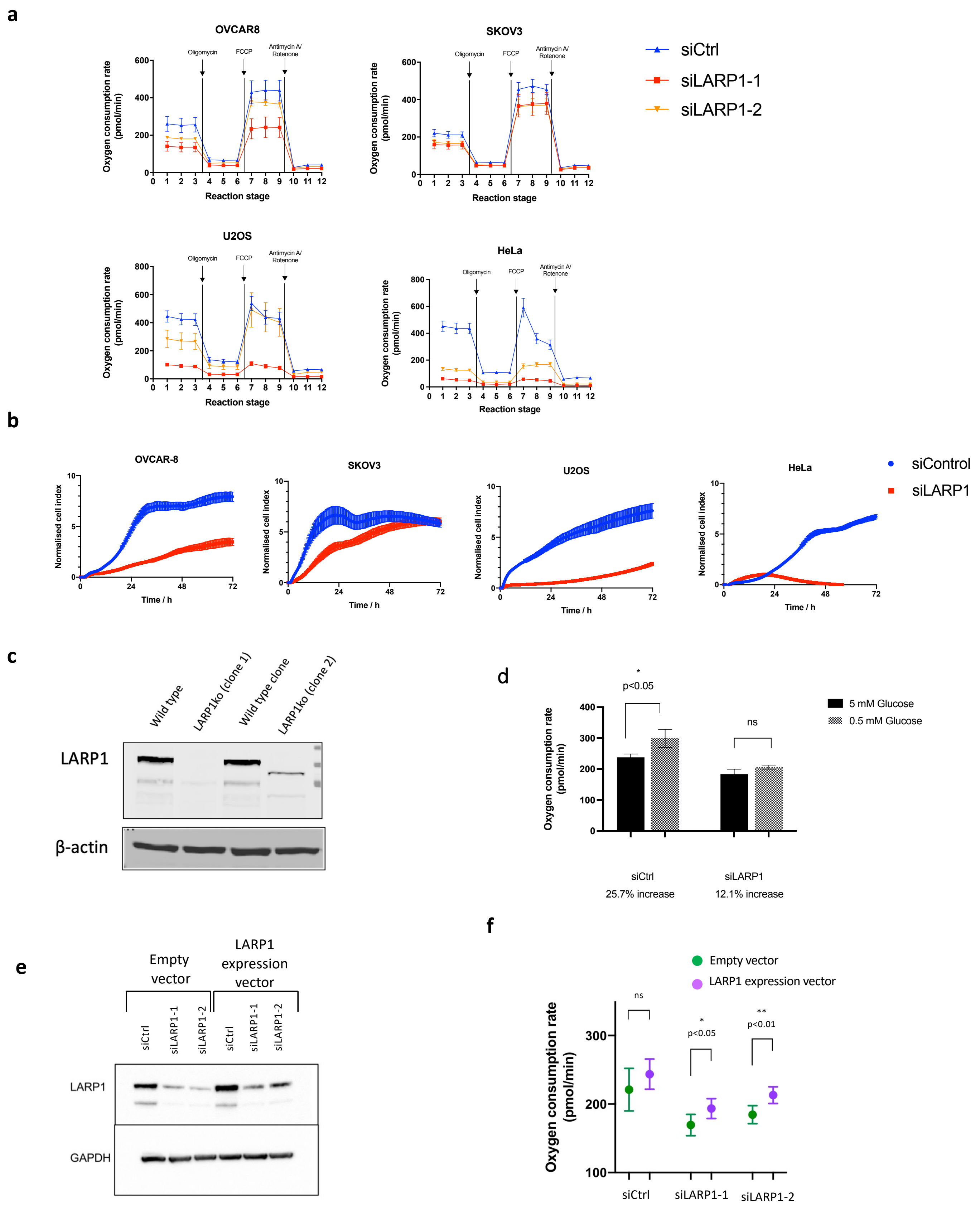

### Supplementary Figure S4

Control siRNA

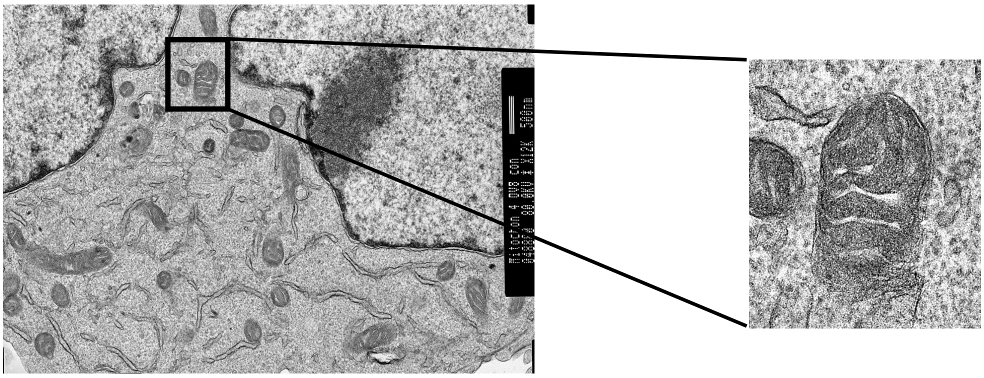

LARP1 siRNA

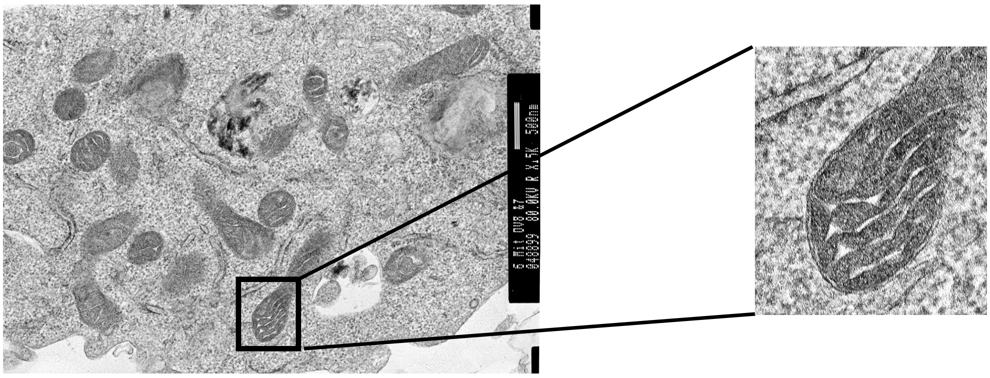

#### Supplementary Figure S5

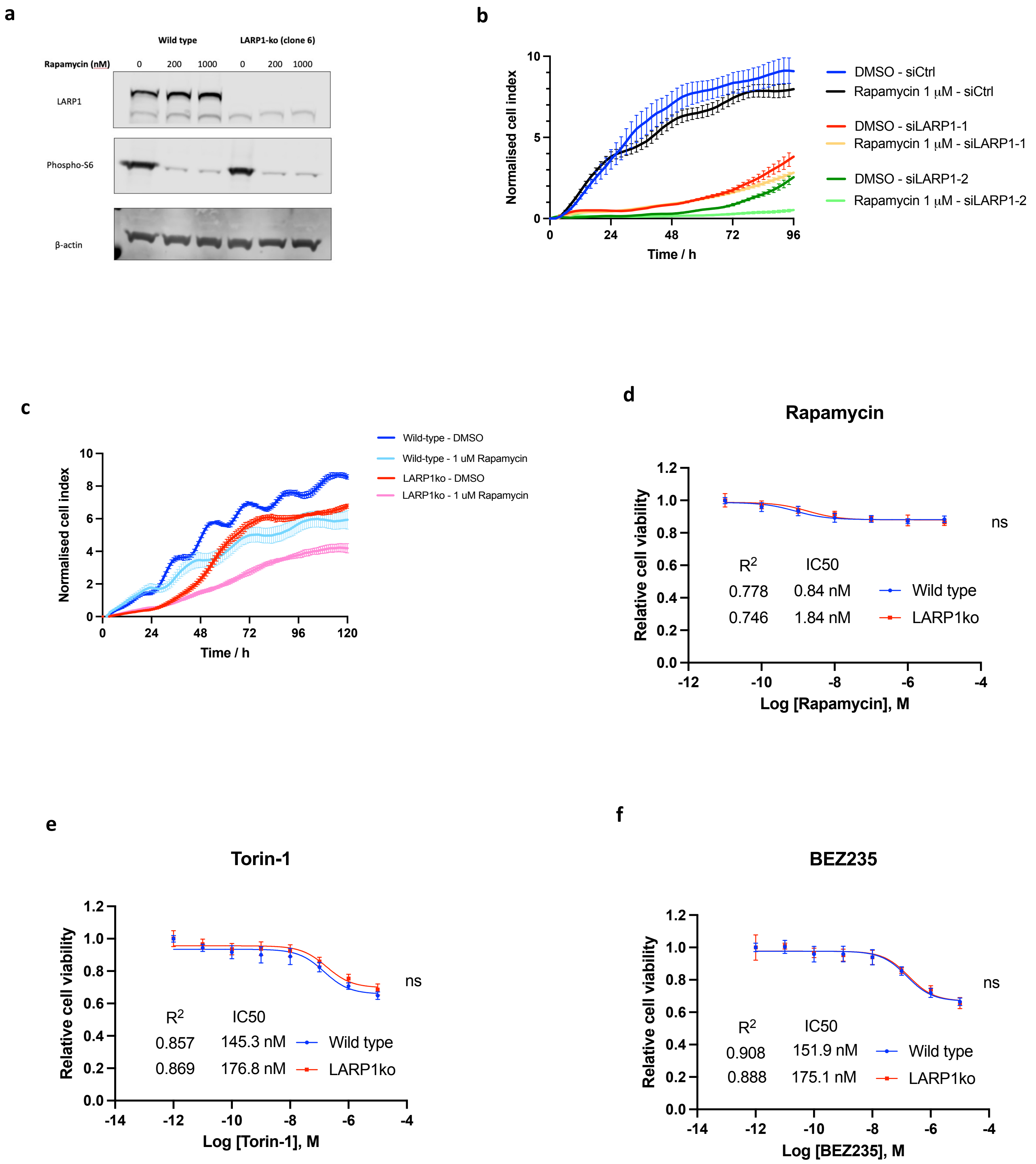

#### Supplementary Figure S6

a

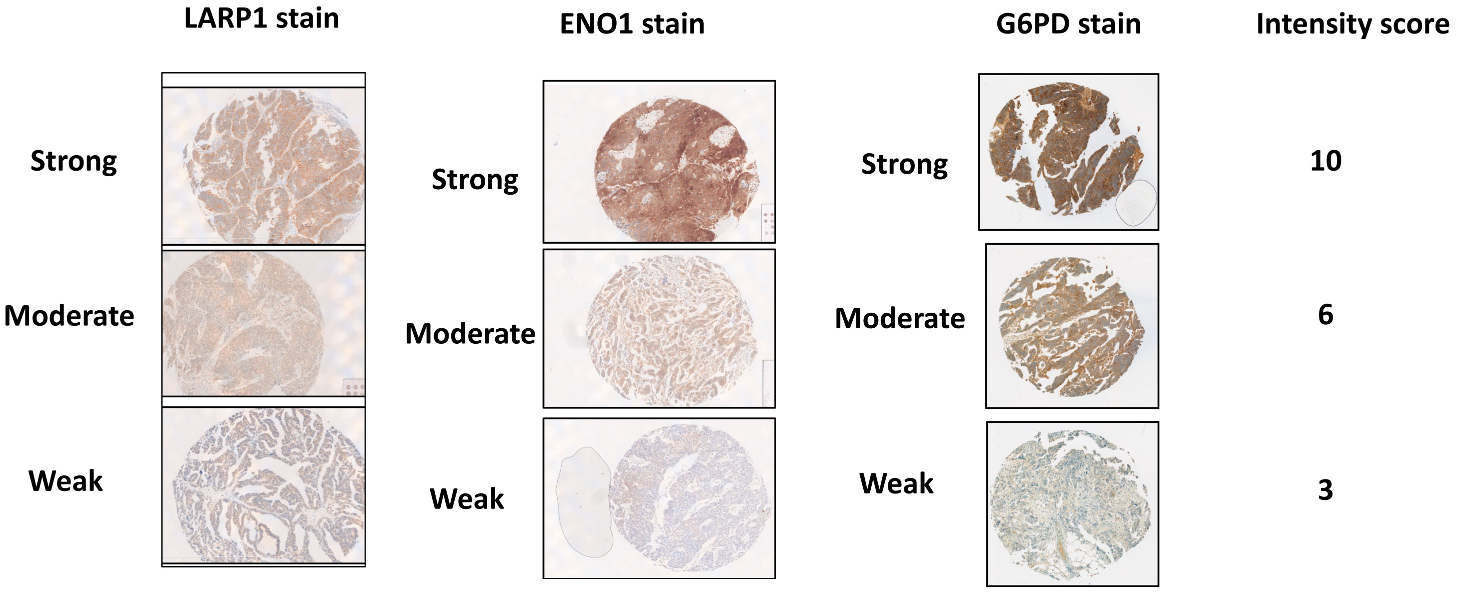


b

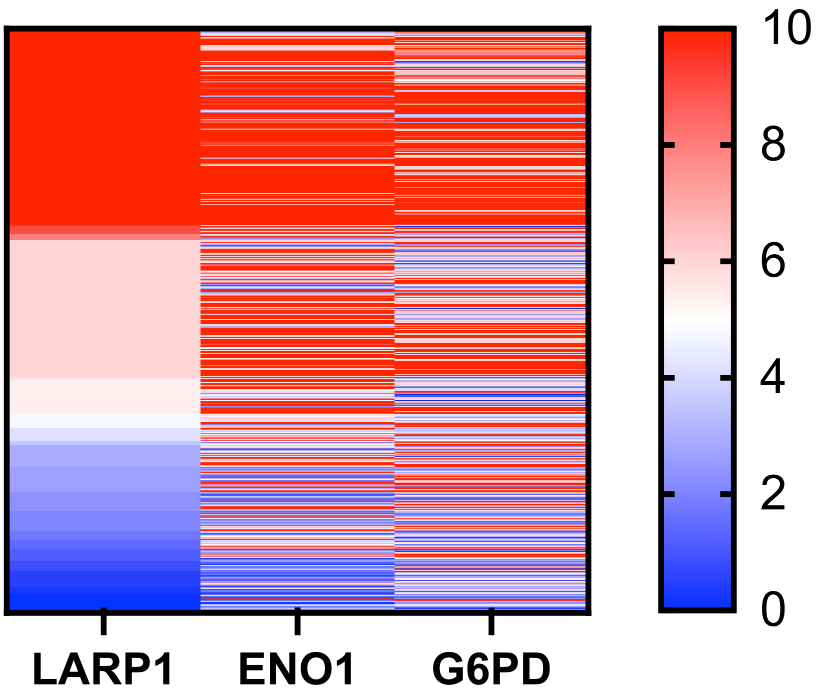
