## Supplementary Figure legends for "LARP1 regulates metabolism and mTORC1 activity in cancer"

**Supplementary Figure S1.** (a) Confirmation of LARP1 knockdown in protein lysates used in proteomics LC-MS analysis. (b) Immunoprecipitation of LARP1 with either anti-LARP1 antibody or IgG isotype control antibody in OVCAR-8 and HeLa cells. Input lysates are also shown. (c) Electrohoretic mobility shift assay for full-length LARP1 or DM15 domain binding to *G6PD* mRNA.

**Supplementary Figure S2.** Metabolite changes in pyrimidine and purine metabolism pathways induced by LARP1 knockdown.

**Supplementary Figure S3** (a) Full oxygen consumption rate traces of MitoStress Seahorse assays followingLARP1 knockdown in OVCAR-8, SKOV3, U2OS and HeLa cells. Basal oxygen consumption rates are plotted in Main Figure 1b. (b) Proliferation rates as measured by Xcelligence real-time cell viability assay following LARP1 knockdown across four different cell lines. (c) Western blots showing knockout of LARP1 by CRISPR-Cas9 in OVCAR-8 clones generated by CRISPR-Cas9 and monoclonal selection. (d)Oxygen consumption rates in OVCAR-8 cells 48 h after LARP1 knockdown, measured either in media containing 5 mM glucose or 0.5 mM glucose (**e,f**) pcDNA4-LARP1 expression vector was transfected into OVCAR-8 cells simultaneously with siRNAs targeting LARP1 to determine if the expression vector could rescue the knockdown phenotype. (e) western blot of LARP1 expression 48 h after co-transfection of the expression vector with siRNAs (f) Oxygen consumption rates measured 48 h after co-transfection of the expression vector with siRNAs.

**Supplementary Figure S4.**. OVCAR-8 cells were transfected with a control siRNA or an siRNA targeting LARP1 and imaged by electron microscopy 48 h after transfection. No effect on mitochondrial morphology was observed. Representative of nine different images taken across three independent samples per group.

**Supplementary Figure S5**.
(a) Wild type or LARP1-ko OVCAR-8 cells were treated with rapamycin at 200 nM or 1 μM to inhibit mTORC1. mTORC1 inhibition was confirmed by dephosphorylation of ribsomal protein S6. Blots were normalised with β-actin. (b) Proliferation rates of OVCAR-8 cells pre-treated with rapamycin and then transfected with LARP1 siRNAs was measured by Xcelligence (c)Proliferation rates of wild-type or LARP1 knockout OVCAR-8 cells pre-treated with rapamycin was measured by Xcelligence
(**d-f**) Inhibition of cell viability with different concentrations of the mTOR inhibitors rapamycin, Torin-1 and BEZ235 were measured in wild-type and LARP1ko OVCAR-8 cells to calculate the $IC_{50}$ values of these drugs. Pairwise comparisons performed using Kolmogorov-Smirnov test *, ($p<0.05$), ** ($p<0.01$).

**Supplementary Figure S6**
(a)Scoring chart for intensity of LARP1,ENO1 and G6PD staining of ovarian cancer tissue microarrray (TMA). Cores were marked and scored as strong (10), medium (6), weak (3) or absent (0) as shown. Intensity score was multiplied by coverage score to yield a combined score for each core. (b) Heatmap showing all cores on the TMA ranked by LARP1 score. Each

core is represented by a horizontal line on the heatmap showing its LARP1, ENO1 and G6PD score by colour.

### SUPPLEMENTARY TABLES

**Table S1. Results of Proteomics LC-MS analysis.** Relative levels of individual proteins in LARP1 knockdown cell lysates compared with control samples.

**Table S2. The LARP1 interactome in HeLa cells as previously described in Mura et al., 2015.**

**Table S3. Analysis for significantly enriched GO-BP pathways within the LARP1 interactome**. Metabolic/catabolic-related processes are highlighted in blue, biosynthetic processes in green. LARP1 interactome is defined as in Table S2.

**Table S4.** Primer sequences used in RT-qPCR

**Table S5.** Antibodies used in western blotting

**Table S6.** Oligo probe sequences used in electrophoretic mobility shift assays
